# Tracking self-citations in academic publishing

**DOI:** 10.1101/2019.12.20.884031

**Authors:** Ameni Kacem, Justin W. Flatt, Philipp Mayr

## Abstract

Citation metrics have value because they aim to make scientific assessment a level playing field, but urgent transparency-based adjustments are necessary to ensure that measurements yield the most accurate picture of impact and excellence. One problematic area is the handling of self-citations, which are either excluded or inappropriately accounted for when using bibliometric indicators for research evaluation. Here, in favor of openly tracking self-citations we report on self-referencing behavior among various academic disciplines as captured by the curated Clarivate Analytics Web of Science database. Specifically, we examined the behavior of 385,616 authors grouped into 15 subject areas like Biology, Chemistry, Science & Technology, Engineering, and Physics. These authors have published 3,240,973 papers that have accumulated 90,806,462 citations, roughly five percent of which are self-citations. Up until now, very little is known about the buildup of self-citations at the author-level and in field-specific contexts. Our view is that hiding self-citation data is indefensible and needlessly confuses any attempts to understand the bibliometric impact of one’s work. Instead we urge academics to embrace visibility of citation data in a community of peers, which relies on nuance and openness rather than curated scorekeeping.

## Introduction

Metrics of productivity can be valuable in assisting evaluation, but to do this they must provide complete and accurate descriptions of citations (Cousijn et al., 2018). Currently this is not the case. Fixing the problem is deceptively simple for a variety of reasons, one being that there is no consensus on how to handle self-citation data. The ongoing debate is contentious and further complicated by the widespread use of the *h*-index for research evaluation, which as the dominating metric puts the emphasis squarely on citations to guide decision making (Hirsch, 2005; Hicks et al., 2015). Without question, this creates a real career motivation to strategically use self-citation (Seeber et al., 2019), but this does not in any way diminish the value of self-cites that result from productive, sustained, leading-edge efforts (Cooke and Donaldson, 2014). When used appropriately, self-cites are equally important as cites from the surrounding community, and without tracking them it is impossible to see how scholars build on their own work.

Despite this, many favor a curated form of the *h*-index as a response to the gaming problem. Curation involves hacking away at the citation data to neatly remove all occurrences of self-citation. While such treatment effectively silences direct attempts to boost citation scores, it does not prevent indirect manipulation and also produces undesired side effects. For example, curation ignores when authors use self-citation to attract cites from others, which is alarming given that each self-citation appears to increase the number of citations from others by about one after a year, and by about three after five years (Fowler and Aksnes, 2007). Furthermore, curation unfairly punishes good citation practices, a particularly worrisome issue for those publishing novel ideas or results that challenge well-established dogma. In such cases, self-citation data can be critical as paper outputs may require a substantially longer period of time to attract the attention (i.e., citations) they ultimately deserve. Thus it is not good practice to hide self-citation data. The end result is a distorted record of progress and discovery.

The sensible alternative to curated scorekeeping would be to consider all citation data including self-citations. Towards this goal, we demonstrate an easy way to track self-cites without distorting other metrics, namely the *h*-index. The approach is not meant to criminalize self-referencing, nor do we intend to suggest a certain threshold of acceptable behavior like what Hirsch did when proposing the *h*-index (Hirsch, 2005). Rather we see this as a tool to clarify how researchers build on their own ideas, as well as how self-citing contributes to the bibliometric impact of their own work. Furthermore, researchers are less likely to blatantly boost their own citation scores (Zhivotovsky and Krutovsky, 2008; Bartneck and Kokkelmans, 2011) while others are watching.

### Defining and tracking self-citation in academic publishing

For author-level tracking, we define a self-citation as any instance where a given author cites their own articles. How we define self-citation differs from recent work done by Ioannidis et al. (2019) where they count a self-citation as any occasion where an author of a given article cites that article. Our reason for this is that we want to know how often specific authors self-cite, not how often an article gets cited by coauthors. In general, we believe that authors’ citations should be sorted by source for clarification: self, nonself, coauthor, etc. and tracked separately. We focus here on self-citation data to show how the approach could work.

To study self-citation practices and regularities in field-specific contexts we analyzed citation data from the 2016 Web of Science database via the German Competence Centre for Bibliometrics. Authors with a unique identifier (either ORCID or ResearcherID) were classified into research areas based on the Web of Science categorization scheme. At the level of category, we found that self-citation accounts for only a small percentage of total citations (Figure 1A), however, there is high variation in self-referencing patterns among individual authors (Figure 1B). This heterogeneity has implications for how we make generalizations regarding self-citation practices. In particular, it does not make sense to use summary statistics to capture average trends when averages would provide such a poor description of individual behavior. There is no metric substitute for expert peer review.

**Figure 1.**
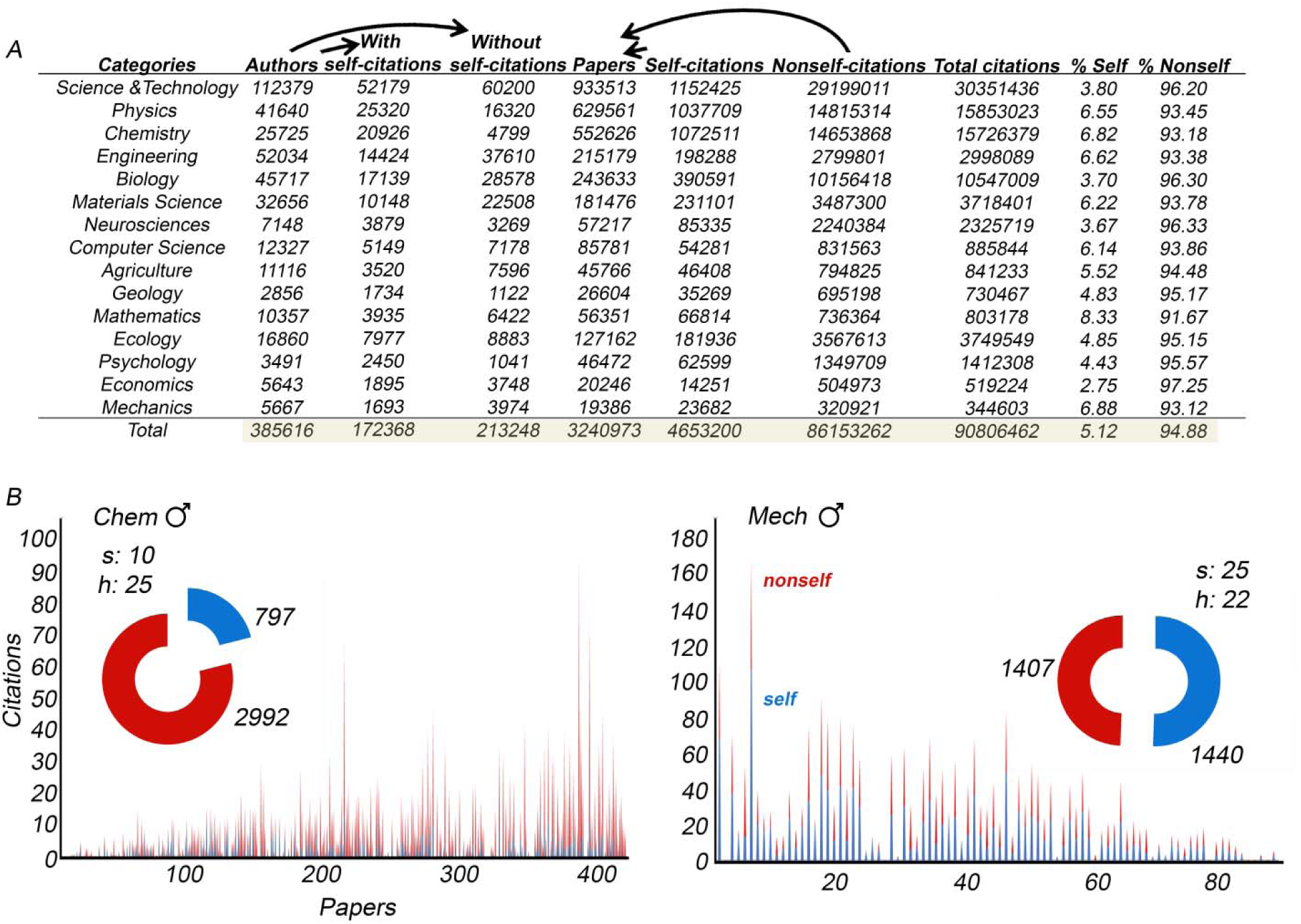
Tracking authors’ self-citation. A) Authors were categorized into 15 subject areas and citations were sorted based on whether they were self or nonself in origin. The total amount of authors, papers, and citations analyzed is highlighted in brown. B) Examples of author profiles where citations have been clarified to separate self- (blue) and nonself-citations (red). Papers are sorted from oldest to most recent. Gender (manually verified), category, s-index, and h-index (excluding self-citations) are reported.

### Computing *s* alongside *h*

We recently proposed a self-citation index (*s*-index), which is little more than a modified

*h*-index.

A scientist has index *s* if *s* of his or her *N*_*p*_ papers have at least *s* self-citations each and the other (*N*_*p*_ -*s*) papers have ≤ *s* self-citations each (Flatt et al., 2017).

The rationale behind the *s*-index is that it brings urgently needed context (e.g., self-citation data) to the *h*-index without introducing distortions. This is especially important given that the *h*-index has become immovable for evaluation and comparison purposes. Towards implementation, we have computed *s*-index scores for all the authors in our study (Figure 2A). From the reported scores, it can be seen that 98% (377,533) of authors achieve an *s*-index of 5 or less, 1.9% (7,561) achieve a score between 6 and 10, and only 0.1% (522) exceed 10. In terms of high scorers, the research categories most represented were Chemistry (236), Science & Technology (97), Physics (73), and Biology (55) (Figure 2B).

**Figure 2.**
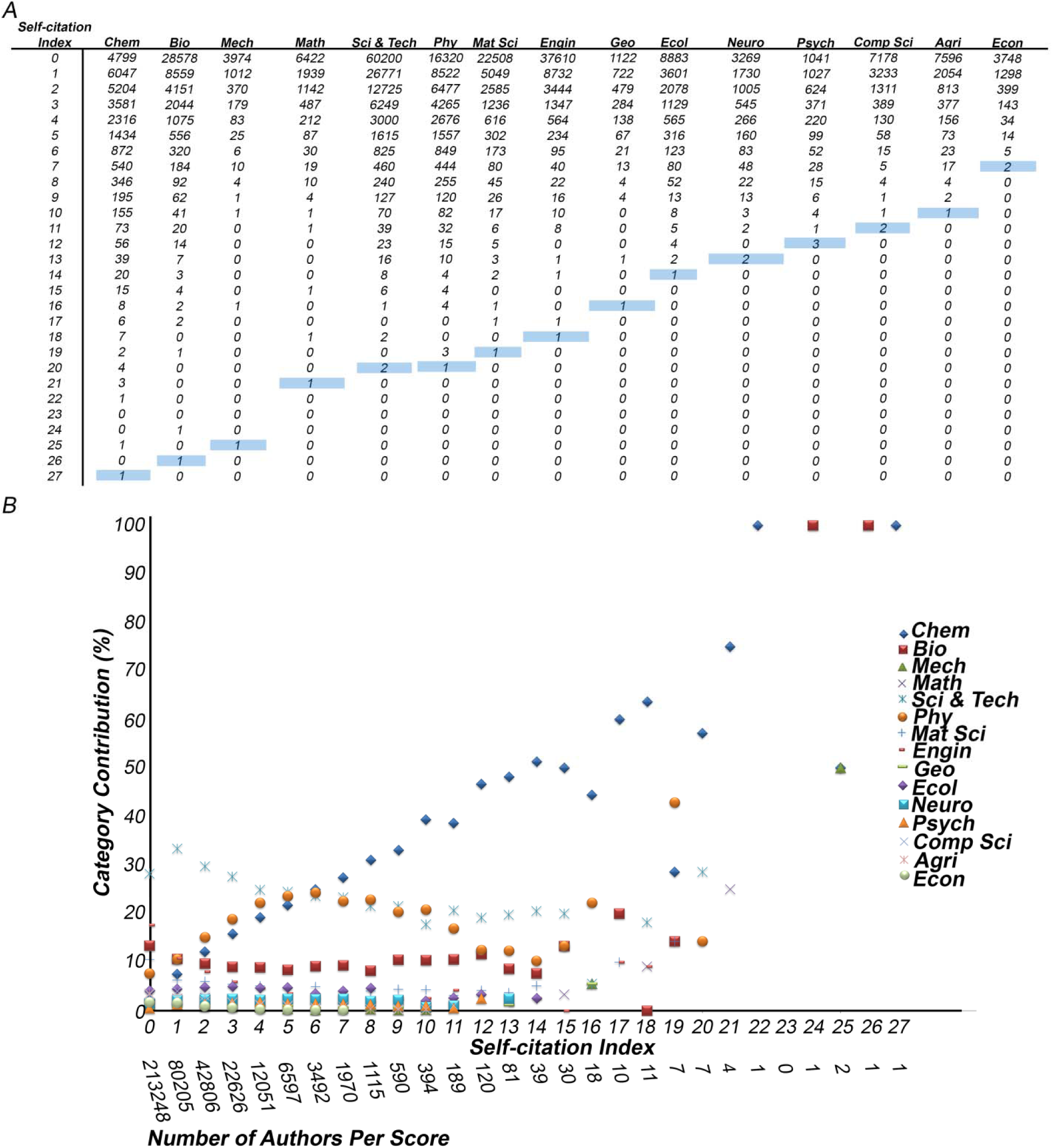
*S*-index scores in field-specific contexts. A) Table shows how authors are distributed according to *s*-index for the 15 categories considered in this study. Highest scores are highlighted in blue (e.g. Chem – 27, Ecol – 14). B) Graph depicts how the categories contribute to observed *s* outcomes. For example, the category Chem is 100% responsible for the one author that achieves a score of 27, whereas both Chem and Mech contain authors with an *s* of 25, and thus they each contribute 50% to the total (here, 2 authors achieve *s* of 25).

We have also included information about the top *s*-index scorers for the different categories (Figure 3). Each of the authors depicted is male, beyond the “early” phase of their careers, and productive in terms of paper outputs and citations. The key difference being the percentage of total citations that are self-citations, which varies from 3% (Psychology) to 58% (Engineering). This difference only becomes apparent by looking at the ratio of self-citations to total citations, which is hidden if you only consider the *s*-index by itself. Thus, it is important to keep track of all the relevant self-citation information and include it alongside nonself-citation data.

**Figure 3.**
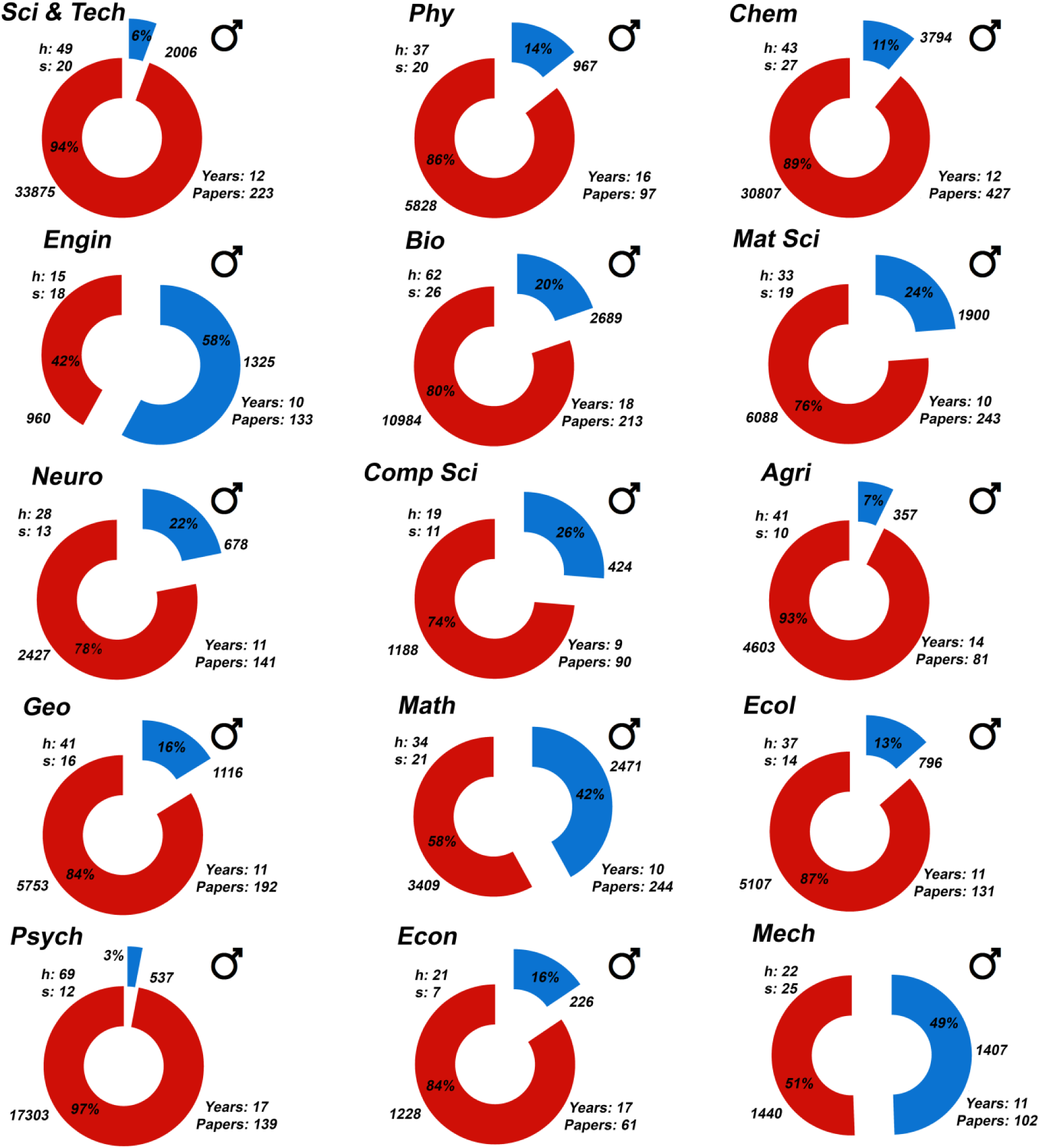
Top *s*-index scorer per category. The highest scorers are depicted along with key information including category, gender (manually verified), *h* (excluding self-citations), *s*, papers, years publishing, and proportion of citations that are self (self-citations in blue and nonself-citations in red).

### Measuring the ratio of self/total citation

Measuring the ratio of self to total citations for authors and each of their papers (see for example Figure 1B) clearly shows the rate at which individuals build on their own ideas in relation to their peers, which is missing in current evaluation procedures. Furthermore, the ratio can serve as an indicator for potential abuse of self-citation by flagging authors that self-cite disproportionately/excessively relative to the norm. In the current study 75% of the authors that self-cite have a ratio below 0.1, meaning that less than 10% of their total citations are self-citations. To see how the measurements breakdown according to category, see Figure 4. Interestingly, there are a total of 1,822 authors in the dataset with ratios that exceed 0.5. Setting thresholds to define acceptable behavior will not explain how such high self/total citation ratios are achievable. Also, doing so will penalize certain situations where a high ratio is legitimate. Rather experts with suitable research backgrounds must help on a case-by-case basis to determine the various factors (e.g., career stage, level of productivity, research topic, citation mentality) that contribute to such high outcomes.

**Figure 4.**
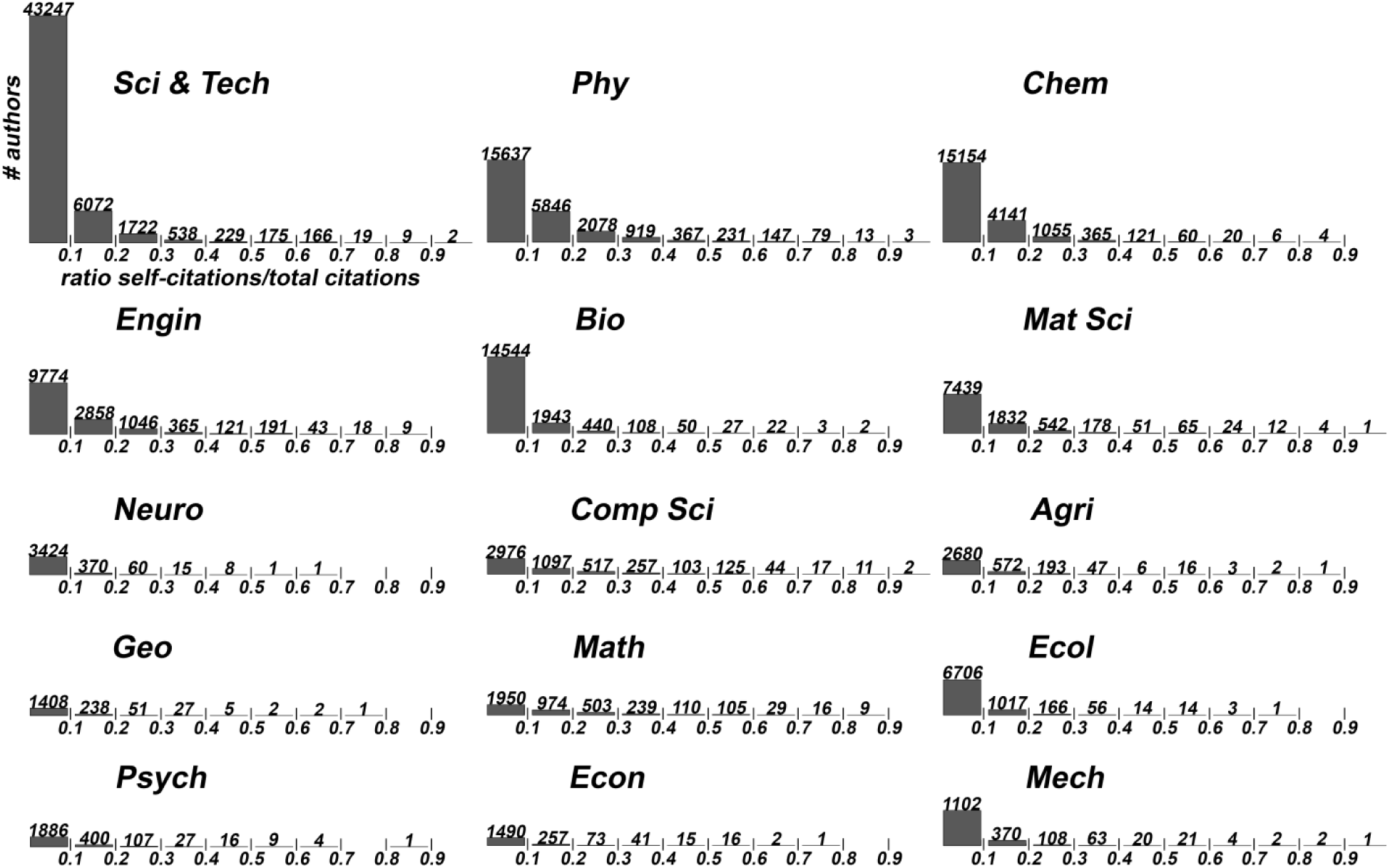
Measuring the amount of self-citation in the total. Self/total citation ratios were calculated for the authors in each category. The exact amount of authors at a particular range of ratio is specified. For example, 15,637 researchers in category Phy have a ratio of self/total citations at or below 0.1.

Figure 5 shows that the ratio of self/total citations tends to decrease as the *s*-index becomes larger, which we observed irrespective of category. Thus in most cases, authors with high *s*-scores are highly cited by their community of peers. This reinforces the idea that we should treat self-citation as a sign of progress (Mishra et al., 2018) rather than viewing the practice suspiciously, or worse scrapping the data altogether. As Cooke and Donaldson have argued, such an observation should be expected from researchers that have published year after year on focused topics with papers building on their own ideas and discoveries (Cooke and Donaldson, 2014). It does not make sense to hide self-citation data when making bibliometric assessments. And importantly, when concerns arise over behavior, for example, when an author has a high ratio of self/total citations along with a high *s*-index, expert panels will have all the relevant data to aid in making sound judgments.

**Figure 5.**
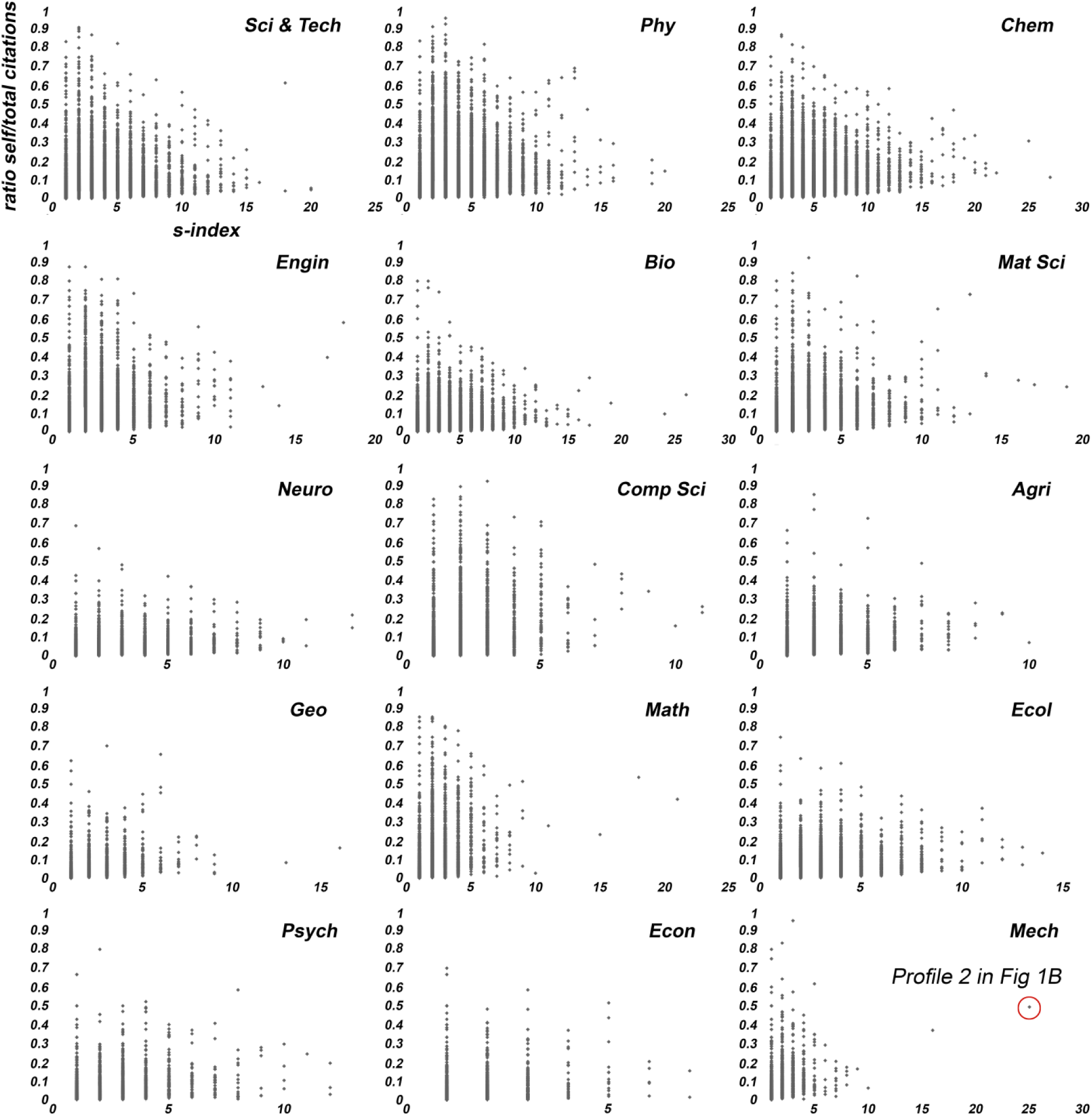
The ratio of self/total citation as a function of increasing *s*-index.

### Conclusion

Clarifying citation buildup will help to accurately evaluate how the various author-related factors influence bibliometric footprint. The justification for doing this is that large-scale citation networks are heterogeneous and contain diverse citation mixing patterns that cannot be summarized using a single global measure (e.g., total citation counts, *h*-index). Instead of curation, we should adopt methods that utilize all the citation data, but in a way that carefully accounts for factors such as self-citation, collaboration, and “citation farms”. Only then can we begin to fully appreciate authors’ behavior and performance in relation to citation records. Towards this goal, we have shown here how to account for self-citation without introducing distortions. Future efforts centered on clarifying citations will better inform policymakers, funding agencies, hiring / promotion / award committees, and the general public about the value of published research.

## Materials and Methods

We extracted authors’ citation data from the 2016 Web of Science (WoS) database produced by Clarivate Analytics and hosted by the Leibniz Institute for Information Infrastructure (FIZ Karlsruhe). The database contains 50,040,717 records for a period of publishing from 1965-2016. Only authors possessing a unique identifier, either ORCID or ResearcherID, were included in the study. For sorting authors into specific research domains we utilized the WoS subject categorization scheme, where we define a given author’s primary field of study as the category (e.g., Physics, Chemistry, Biology) containing the highest volume of publications. We analyzed categories if they contained at least 1000 unique authors. Citation queries were run using the Oracle SQL language.

## Acknowledgements

This work was funded by BMBF (Federal Ministry of Education and Research, Germany) under grant number 01PQ17001, the “Analyzing Self-citations in Web of Science (WoS)” project (http://bit.ly/Selfcitations-project), in the Competence Centre for Bibliometrics.

